# SilencioM: silencing connected interfering oligonucleotides against *MAPT* as a potential disease modifying therapeutic for tauopathies

**DOI:** 10.64898/2026.11.09.765484

**Authors:** Paraskevi Sgourdou, Nicholas Vrettos, Kevt’her Hoxha, Thallyah Baffic, Edward B. Lee, Hong Xu, Virginia M-Y Lee, Zissimos Mourelatos

## Abstract

Tauopathies are a diverse group of neurodegenerative diseases whose pathogenesis involves brain accumulation of misfolded, pathological tau (Microtubule Associated Protein Tau –MAPT–). More than twenty tauopathies have been identified thus far that collectively impact millions of people worldwide, of which the most common is Alzheimer’s disease (AD). Despite substantial efforts to develop drugs for tauopathies, no effective treatments currently exist, highlighting the urgent need for therapeutics to combat these prevalent and ruinous neurodegenerative diseases. Strategies that target tau are likely to have the greatest impact since tau is the proximal cause of their pathogenesis. Small interfering RNAs (siRNAs) have emerged as a powerful class of precision therapeutics that silence disease-causing genes through RNA interference with favorable safety profiles. Delivery challenges are rapidly being addressed, and siRNA-based approaches are poised to transform the therapeutic landscape of neurological diseases. In this study, we describe SilencioM (<u>Silen</u>cing <u>c</u>onnected interfering <u>o</u>ligonucleotides against *<u>M</u>APT*) and show its efficacy in markedly reducing tau pathology after a single intracerebroventricular injection in PS19 mice, a mouse model of severe tauopathy. SilencioM targets human, primate, rat and mouse ortholog genes; thus, the same compound can proceed from efficacy studies to Investigational New Drug Application (IND)-enabling studies. SilencioM is simple to manufacture and is administered without encapsulation in lipid nanoparticles and without requirements for lipophilic conjugation, making it a promising candidate for further development as a novel therapeutic for tauopathies, including AD.

## INTRODUCTION

Tau, encoded by the Microtubule-Associated Protein Tau (*MAPT)* gene, plays an important role both in maintaining overall neuronal structure by stabilizing microtubules and establishing cell polarity, as well as regulating axonal transport [20, 68, 74]. Alternative splicing of *MAPT* messenger RNA (mRNA) generates six tau isoforms, differing by the presence of the number of N-terminal repeats (N0, N1, N2) and the presence of either three (3R) or four (4R) microtubule-binding repeat domains [8] [68]. Misfolding of tau into amyloid fibrils (filaments) [61] in specific brain regions results in impaired synaptic signaling, disrupted neuronal function, and ultimately neuronal death, and underlies the pathogenesis of a large class of neurodegenerative diseases known as tauopathies [48] [46] [24]. Pathological tau fibrils form intracellular aggregates known as neurofibrillary tangles (NFTs), pre-tangles and neuropil threads (NTs), and these aggregates have the capacity of spreading along anatomically connected pathways, seeding the conversion of soluble tau into more pathological tau, which leads to disease progression, neuronal loss and degeneration [60] [24].

A subset of tauopathies is caused by genetic variants: such as *MAPT* itself, directly disrupting *MAPT* physiology, or *VCP* (valosin-containing protein), indirectly influencing proteostasis [24]. However, most primary tauopathies are sporadic and include diseases such as Progressive Supranuclear Palsy (PSP) and Corticobasal Degeneration (CBD), where aggregation of 4R tau predominates; Pick’s disease (PiD), a 3R tauopathy; and primary age-related tauopathy (PART), a 3R+4R tauopathy [48] [23] [19] [24]. There are many more diseases (secondary tauopathies) whose pathogenesis is associated with the accumulation of pathological tau, the most common of which is Alzheimer’s disease (AD), a 3R+4R tauopathy, that also involves deposition of amyloid-β fibrils, derived from amyloidogenic proteolytic processing of Amyloid Precursor Protein (APP) [37] [12] [17] [24] [41]. Tau undergoes multiple post-translational modifications that regulate its physiological function in microtubule binding and stabilization, as well as its pathological aggregation in tauopathies [4, 75] [24]. Notably, hyperphosphorylation is a hallmark of pathological tau [36] [47] and is thought to be an early pathogenetic event as it promotes aggregation by impairing tau’s microtubule binding and lowering its solubility [13] [3] [2].

Considerable efforts to develop disease-modifying drugs for tauopathies have been undertaken, but there are no therapies yet, underscoring the need to therapeutically target tau. One strategy involves passive immunotherapy, to intercept extracellular misfolded tau transmitted trans-neuronally via extracellular release followed by uptake of pathogenic seeds that promote tau pathology in recipient neurons. After initial failures using antibodies targeting N-terminal tau, newer antibodies are currently being evaluated in clinical trials [64] [22]. A challenge for immunotherapies is that the vast majority of pathological tau is inaccessible to antibodies, as it is intracellular or enclosed in extracellular vesicles during intercellular transmission of tau seeds [78] [73]. Thus, antibodies in the interstitial fluid will only have access to the tiny portion of tau that may be extracellular, secreted as free protein [16]. Furthermore, tau seed heterogeneity complicates effective targeting of all species by monoclonal antibodies that typically recognize a single epitope [21]. An alternative strategy is to limit tau production by reducing *MAPT* mRNA levels using antisense oligonucleotides (ASOs) or small interfering RNAs (siRNAs) [67] [30] [45] [11] [25] [29]. This approach can target the accumulation of both intracellular and extracellular proteins that in the case of tau is crucial for the reduction of all its pathological forms, most of which are found in the cytoplasm and dendrites of neurons. Intrathecally administered ASOs against *MAPT* have advanced from preclinical studies [27] to human clinical trials, with BIIB080/Diranersen (Biogen) currently being tried for AD, and NIO752 (Novartis) for PSP and AD [7]. In the Phase 2 CELIA trial, the intrathecally administered MAPT-targeting ASO diranersen (BIIB080) [55] lowered tau pathology and slowed cognitive decline in early AD, but tau reduction was comparable across all doses tested and the primary endpoint of a Clinical Dementia Rating– Sum of Boxes dose–response was not met (NCT05399888). siRNAs are more potent and specific than ASOs: by mimicking microRNA duplexes, siRNAs assemble with the Ago2 nuclease to form multiple turnover enzymes [50] [54]. Traditionally, siRNAs suffered from poor Central Nervous System biodistribution, but this hurdle is rapidly addressed with new siRNA designs, chemical modifications and conjugates [1] [5] [15] [29] [66]. Modified siRNAs persist in endolysosomes, which act as a reservoir, creating a “depot effect” that enables the prolonged, low-level release of functional siRNA into the cytoplasm over time. This mechanism contributes significantly to the remarkable durability of the therapeutic effect of modified siRNAs [14]. Collectively, these new approaches allow direct administration of siRNAs in Cerebrospinal Fluid (CSF), which leads to widespread, specific, potent and sustained gene silencing in the brain and spinal cord of rodents and primates, without apparent toxicity [5] [15] [66]. A pivotal early study showed that intrahippocampal injection of a mixture of four *MAPT*-targeting siRNAs (Accell™ SMART pool) led to local knockdown and reduction of tau pathology in the P301S mouse model of tauopathy, lending support to the use of siRNAs to target tauopathies [76]. Currently early phase clinical trials are underway for LY3954068 (Eli Lilly), an siRNA targeting *MAPT* delivered intrathecally; and ARO-MAPT (Arrowhead Pharmaceuticals) and DNL-628 (Denali), which are siRNA-brain shuttle conjugates administered systemically for AD and tauopathies [38] [7]. Targeting *MAPT* with siRNAs (ARO-MAPT) led to dose-dependent reduction of tau in CSF without adverse effects in healthy volunteers [6].

In this study, we have developed a <u>Silen</u>cing <u>c</u>onnected interfering <u>o</u>ligonucleotide against *<u>M</u>APT* (SilencioM). We show that a single intracerebroventricular (ICV) injection of SilencioM in PS19 mice, a widely used mouse model of severe tauopathy [77] [69] markedly reduces tau pathology. SilencioM targets human, primate, rat and mouse ortholog genes; thus, the same compound can proceed from efficacy studies to Investigational New Drug Application (IND)-enabling studies. SilencioM is simple to manufacture and is administered without encapsulation in Lipid Nanoparticles (LNPs) and without requirements for lipophilic conjugation, making it a promising candidate for further development as a novel therapeutic for tauopathies, including AD.

## MATERIALS AND METHODS

### Mouse lines

All mouse experiments were approved by the Institutional Animal Care and Use Committee (IACUC) of the University of Pennsylvania and were performed in accordance with the ethical guidelines of the National Institutes of Health (NIH). Mice were raised under a constant 12-hour light/dark cycle. B6 (C57BL6) and PS19 mice (B6.Cg-Tg(Prnp-MAPT*P301S)PS19Vle/J, Strain: 024841) were purchased from Jackson Labs. Both male and female mice were used in these studies.

### Oligonucleotides used for gene silencing

For the initial *in cellulo* screens, siRNAs were synthesized by Integrated DNA Technologies (IDT). Modified oligonucleotides for NTC were synthesized by Gene Link. Preclinical/therapeutic grade modified oligonucleotides for SilencioM were synthesized and purified by Eurogentec. Lyophilized oligonucleotides were resuspended in artificial CSF (aCSF) buffer (137 mM NaCl, 5mM KCl, 20mM D-(+)-Glucose, 8mM HEPES, pH 7.4). To generate SilencioM, two parts of M2024-Guide were annealed to one part of M-passenger by heating at 65^0^ C for 5 minutes, followed by 37^0^ C for 60 minutes and cooled on bench to room temperature. Stock concentration of SiliencioM was at 8mM; dilutions were made with aCSF. To generate NTC, equal parts of NTC-Guide1, NTC-Guide2 and NTC-Passenger, were annealed.

### Stereotactic intracerebroventricular (ICV) injections

The mice underwent bilateral, intracerebroventricular (ICV) injection by insertion of a single needle into the brain at established coordinates from Bregma using a stereotactic rack: +0.2mm to +0.3 mm Anterior, −1.00 mm Lateral (right) and +1.00mm Lateral (Left), −2.5mm to −3.00mm Ventral. Each of the two injection sites received 5 μl of 2 mM SilencioM, or 2 mM NTC (both dissolved in aCSF) or aCSF only. The procedure was as described in [39]. Briefly, mice were deeply anesthetized with ketamine/xylazine/acepromazine, immobilized in a stereotaxic frame (David Kopf Instruments). The injection on either site was performed at a rate of 0.4 μl/min followed by a 2 min. wait before the needle was pulled out. The body temperature of the mice was maintained at 37 °C with a heating pad during surgery while subsequently they were placed under a heating lamp and monitored until awake and exhibiting voluntary locomotion (≥ 2 hours). Mice were also monitored for a minimum of 3 days after surgery.

### Cell culture and transfection

HEK-293T cells and SHSY5Y cells were maintained in Dulbecco’s modified Eagle’s medium (DMEM) and DMEM/F12, respectively, at 37 °C and 5% CO_2_. Media were supplemented with 10% fetal bovine serum (FBS) (ThermoFisher Scientific, Waltham, MA, USA) and 1% Glutamine. Transfections were performed with RNAiMAX Transfection Reagent (ThermoFisher Scientific, Waltham, MA, USA) according to manufacturer’s instructions. The cells were harvested 48h after transfection.

### RNA/protein extraction

Transfected cells were processed with Qiagen RNeasy mini kit for RNA extraction and subsequent reverse transcription-quantitative polymerase chain reaction (RT-qPCR). Frozen brain tissue was ground with plastic pestle in RIPA buffer (50 mM Tris-HCl, pH 7.5, 150 mM NaCl, 5 mM EDTA, 1% NP-40, 0.5% sodium deoxycholate, 0.1% SDS) supplemented with RNAse (1u/ul), protease and phosphatase inhibitors (1X), then sonicated 3 times at 30% output. Half of the sonicated lysate was mixed with Trizol and used for RNA purification with miRNeasy kit (Qiagen). The remaining lysate was centrifuged at 16.000xg for 15 min, at 4°C. The amount of protein in the samples was measured by Bradford assay, and lysates were stored at −80°C.

### mRNA quantification

250 ng of total RNA were reverse transcribed using the Quantitect kit (Qiagen). The produced cDNAs were amplified with PowerUp SYBR-green mix (Applied Biosystems) and the primers used are listed in Supplementary Table 1. Assays were run on StepOnePlus Real-Time PCR system (Applied Biosystems) and quantified with the ΔΔCt method [51] *Gapdh* mRNA was used for input normalization. Results were calculated from 3 technical triplicates and analyzed using Graphpad-Prism 10.4.1.

### Single Molecule Array (Simoa) digital immunoassay

The total amount of human tau protein (expressed by the transgene in PS19 mice) present in each sample lysate was quantified using the ultrasensitive Single Molecule Array (Simoa) digital immunoassay platform on a Quanterix HD-X Analyzer (Simoa® Tau v2.0 Advantage Kit, catalog number 101552). Samples were prepared according to the manufacturer’s protocol, involving centrifugation followed by dilution with kit-specific sample diluent to reach the Minimum Required Dilution (MRD). Each sample, along with the appropriate negative and positive controls, was run in technical duplicates.

### *In Situ* Hybridization (ISH) with Digoxigenin-labeled Locked Nucleic Acid (LNA) probes

The procedure was adopted from [57] with few modifications. Briefly, we designed an LNA probe against M2024 (5’-AG+GTGGCCA+GGT+GGAAG+TAAA, +denotes LNA base) that was synthesized at IDT. The probe was digoxigenin labeled with Dig-ddUTP and the Dig-3’-end labeling kit (Roche) and stored at −20^0^ C until use. 5 μm-thin sections of FFPE (fresh frozen paraffin embedded) tissue were placed onto Probe-on slides. Slides were deparaffinized in xylenes, 2 x 5 min followed by 5 min each in serial dilution of ethanol (100%, 100%, 75%, 50%, 25%), followed by 2 changes of ddH_2_O. Unmasking of tissue RNA was done using Heat induced epitope (antigen) retrieval (HIER) using basic buffer (10 mM Tris-base, 1 mM EDTA solution, pH 9.0) from Vector Labs (H-3301-250). 1X Basic-Hier solution was pre-heated in a glass beaker on a hot plate until temperature reached 95-100 °C. The slides were immersed in staining rack in the heated buffer for 10min (up to 15 min) and then cooled down at RT. Subsequently, slides were digested with proteinase K solution (1X) for 10 min at 37^0^ C, rinsed in PBS (2 rinses) and incubated with Hybridization mix (miRNA ISH buffer and LNA Detection Probes) for 1h at 55°C. Sections were subjected to stringent washes at 55°C in 5xSSC (1time), in 1xSSC (2 times), 0.2xSSC (2 times) and 0.2xSSC (1 time) at RT before transferring to PBS. Sections were blocked for 1hr in blocking solution (2% sheep serum, 2mg/ml BSA in PBST) at RT followed by incubation with a sheep anti-DIG-AP antibody (Roche catalog # 11 093 274 910, dilution 1:1000) at 4°C overnight in a humidified chamber. Next, slides were washed 4 times, 10 min each in PBST, and the AP reaction was achieved by incubating the slides with 1-Step NBT/BCIP plus Suppressor for approximately 2hours at 30°C in the humidifying chamber protected from light. KTBT buffer was used 2 times for 5 min each to stop the reaction. Slides were washed with water, 2 times for 1 min each, and counterstained with Nuclear Fast Red for 1min. After several changes of H_2_O, the sections were dehydrated by passing through a series of alcohols (50%, 75%, 95%, 100%, 100%) and xylenes and coverslipped in Cytoseal.

### Histology and Immunohistochemistry (IHC)

Adult mice were intracardially perfused with PBS, their brains were removed and mid-sagitally transected into two hemispheres. One hemisphere was immediately submerged in a solution of 10% Neutral Buffered Formalin (NBF) for fixation overnight at 4°C, and the second one was dissected in cold PBS, flash frozen in liquid nitrogen or dry ice for biochemical experiments. The fixed hemisphere was further sliced with a mouse brain slicer and processed for embedding in paraffin blocks. Sections were deparaffinized 2 x 5 min in xylene and immersed in descending EtOH series (2 x 100%, 1 x 95%, 1 x 80%, 1 x 70%) 1 min each. Antigen retrieval was done with Formic Acid (88%) for 5 min, then rinsing with water for 5 min. Slides are then immersed in freshly prepared H_2_O_2_/water (50 mL H_2_O_2_ to 150 mL water/boat) for 30 min followed by wash under running tap water for 10 min and in 0.1M Tris pH 7.6 for another 5 min. Blocking is done with 0.1 M Tris/2% FBS (PH=7.6 needs to be confirmed before use) for 5 min. followed by incubation in Ab solution (Phospho-Tau Ser202, Thr205, AT8 monoclonal: Thermo Fisher Scientific MN1020, 1:1000, or GFAP: Sigma G9269, 1:2000) overnight at 4°C in a humidified chamber. After washing in 0.1M Tris pH 7.6 for 5 min, we block in 0.1 M Tris/2% FBS for another 5 min before incubating in secondary biotinylated anti-mouse antibody 1:1,000 in 0.1 M Tris/2% FBS for 1 hr. at RT. We repeat washing and blocking for 5 min each, followed by incubation in AB solution for 1 h at room temperature. The sections are incubated in DAB, 8min for AT8 and 1.5 min for GFAP (time is kept consistent in all stainings), followed by washes in 0.1 M Tris pH 7.6 for 2 x 2 min and counterstained with Harris Hematoxylin solution for 20 seconds. After washing the sections in running tap water for 10 min we dehydrate in ascending series of ethanol (1 x 70%, 1 x 80%, 1 x 95%, 2 x 100%), 1 min each, and clear in 2 changes of xylene, 5 min each, before coverslipping in Cytoseal. The slides were scanned with 3DHistech Pannoramic 250 while AT8 and GFAP expression was quantified with QuPath software, version 0.6.0 [9].

### Statistical analysis

Statistical analysis was performed using unpaired one-tailed Welch’s t-test, or unpaired Mann-Whitney one-tailed test, depending on whether the data followed normal (Gaussian) distribution or not, respectively. A one-tailed test was specified *a priori* because SilencioM is designed to lower mRNA levels of *MAPT* so only a reduction relative to vehicle (aCSF) was of interest. Analyses and graphing were performed with Graphpad-Prism 10.4.1. The Z-factor [80] calculated for *MAPT* mRNA RT-qPCR assay is 0.85. Data in the graphs are represented as mean ± Standard Error of the Mean (SEM).

## RESULTS

### Design and generation of SilencioM

To target *MAPT* mRNA we sought to employ siRNAs [30], as they emulate the endogenous miRNA pathway and assemble with the Ago2 nuclease to form multiple turnover enzymes that potently silence target genes [40] [54] [50] (Figure 1a). We first designed individual siRNAs against *MAPT* using the following criteria: a) optimal design principles that promote loading of the guide strand to Ago2, as well as target accessibility and specificity (reviewed in [70]); b) the ability to target and silence all human *MAPT* mRNA isoforms; c) an ability to target *MAPT* homologs of multiple species: human, mouse, rat, cynomolgus macaque; and d) the ability to target in addition to the wild-type human *MAPT* allele, all major disease-associated variants. We tested their efficacy in reducing *MAPT* mRNA *in cellulo,* using human 293T cells, which endogenously express many neuronal genes including *MAPT* [63] [49], and in SH-SY5Y human neuroblastoma cells [10]. We chose M2024 for further development as fully modified duplexes that we refer to as small interfering oligonucleotides (siOs), since all 2’-hydroxyl groups are substituted with either 2’-Fluoro or 2’-O-methyl groups. These modifications are essential for stability and *in vivo* efficacy while also reducing innate immune activation without abrogating Ago loading or silencing potency [1] [18] [44] [62] [29]. We then covalently linked the passenger (sense) strands of siOs, in a 5’ to 3’ manner, by using polyethylene glycol linker, and annealed the guide (antisense) strands to generate SilencioM (Figure 1b). An added benefit of linking the passenger strands is that it renders them long, disfavoring loading to Ago2 and reducing any off-target effects from the passenger strand. The guide strands contain 5΄-(E)-vinylphosphonate (5’-E-VP), rendering them resistant to phosphatases and promoting Ago2 binding [59] [31]. We unpaired the first two nucleotides between the 5’ end of the guide strand and the passenger strand to decrease 5’-terminal thermodynamic stability, which is expected to facilitate Ago2 loading of the guide strand [72]. Phosphodiester bonds were replaced with phosphorothioates in select positions to increase stability and cellular uptake [28] [5] [25]. The final sequences with modifications used in the production of SilencioM and of non-targeting control (NTC) are shown in Table 1. All chemical modifications introduced (Figure 1c), except for 5’-E-VP, have previously been used in FDA-approved ASO or siRNAs [29]. The siOs used to generate SilencioM were subjected to Ultra Performance Liquid Chromatography (UPLC) after synthesis and their identity was confirmed by Mass Spectrometry (MS); representative chromatographs and mass spectra are shown in Supplementary Figure 1. To test the potency of SilencioM in downregulating *MAPT* mRNA, we calculated its half maximal inhibitory concentration (IC50) from 8-point dose-response curves using human 293 cells and found that it is highly potent *in cellulo,* with an IC50 of 2 pM (Figure 1d).

**Figure 1.**
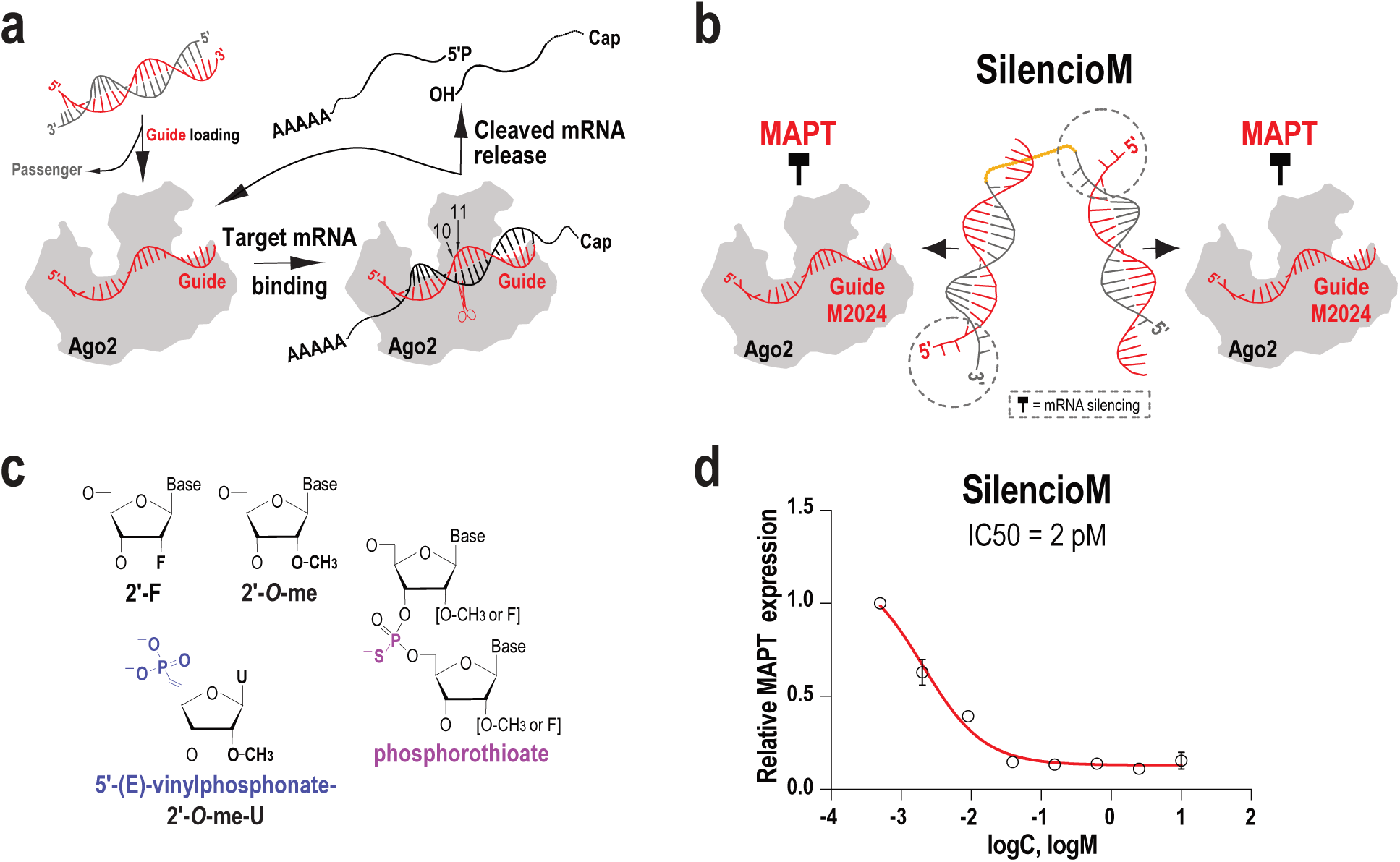
Design and properties of SilencioM. **(a)** Schematic of RNA silencing by Ago2 and siRNAs. The guide (antisense) strand of an siRNA duplex is loaded to the Ago2 protein while the passenger (sense) strand is released. Guide RNA recognizes its target mRNA with antisense complementarity, triggering the Ago2 nuclease domain to cleave the phosphodiester bond of the target mRNA at a position across the 10th and 11th nucleotide of the guide RNA (scissors). The cleaved mRNA is released and further degraded by cellular exonucleases. The Ago2-Guide ribonucleoprotein is then available for further rounds of target mRNA recognition and cleavage, leading to potent gene silencing. **(b)** Schematic of SilencioM: the linker that covalently connects the 5’ end of one passenger strand to the 3’ end of another passenger strand, is shown in orange. Dashed circles show the two unpaired bases between guide and passenger strand. The duplex is generated by annealing 2 parts of M2024-Guide (red) to one part of M-passenger strand (grey). **(c)** Schematic of the modifications used. **(d)** Half-maximal inhibitory concentration (IC50) of SilencioM, in human 293 cells, calculated from 8-point dose response curves performed in triplicates (10 nM, 2.5 nM, 0.625 nM, 0.156nM, 0.039 nM, 0.009 nM, 0.002 nM, 0.0006 nM).

**Table 1.** Sequences and modification of oligonucleotides used for SilencioM and for Non-targeting Control (NTC)

|  |  |
|---|---|
| **M2024-Guide:** | vPmU*fU*mUfAfCfUmUfCmCfAmCfCmUfG*mG*fC*mC*mA*mC*mC*mU |
| **M-Passenger:** | mG*mC*mCfAmGmGmUfGmGmAmAmGmUfA*mU*mU-L36-mG*mC*mCfAmGmGmUfGmGmAmAmGmUfA*mU*mU |
| **SilencioM:** | M2024-Guide + M-Passenger, in 2:1 ratio |
| **NTC-Guide1:** | vPmU*fU*mAfGfUfCmGfAmCfAmUfGmUfA*mA*fA*mC*mC*mA*mU*mU |
| **NTC-Guide2:** | vPmU*fC*mAfCfAfCmAfAmCfAmUfGmUfA*mA*fA*mC*mC*mA*mU*mU |
| **NTC-Passenger:** | mU*mU*mUfAmCmAmUfGmUmCmGmAmCfU*mA*mA-L18-mU*mU*mUfAmCmAmUfGmUmUmGmUmGfU*mG*mA |
| **NTC:** | NTC-Guide1 + NTC-Guide2 +NTC-Passenger, in 1:1:1 ratio |

(m = 2'-O-me;  f = 2'F; vPmU = Uridine with 5' E-vinylphosphonate and 2'-O-me; \*= phosphorothioate; L36, L18 =polyethylene glycol linker containing 36 or 18 atoms, respectively.)

### SilencioM shows widespread distribution and retention in the mouse brain

To test the efficacy of SilencioM in ameliorating tau pathology, we used the PS19 transgenic mouse line (B6;C3-Tg(Prnp-MAPT*P301S)), a widely used animal model that recapitulates key pathological changes seen in human tauopathies [69, 77, 79]. PS19 mice express the T34 isoform (1N4R) of human *MAPT* with a P301S pathogenic variant that causes early-onset familial frontotemporal dementia (FTD), driven by the mouse prion protein promoter (Prnp) [77]. The mutant tau transgene in PS19 mice is expressed at ∼5-fold higher levels than endogenous mouse tau [77].

In initials experiments we performed intracerebroventricular (ICV) injections of 20 nmoles of SilencioM or 20 nmoles of Nontargeting Control (NTC) Silencio, or buffer only in B6 wild-type mice or in PS19 mice, to examine the distribution of SilencioM in mouse brain. In one experiment we injected SilencioM labeled with a red fluorophore (Cy3 conjugated to the 5’end of the passenger strand) or PBS to B6 mice, sacrificed them after 48 hours and macroscopically detected SilencioM, which was distributed throughout the mouse brain (representative image shown in Figure 2a). Next, we ICV-injected a cohort of PS19 mice (5-6.5 months old) with 20 nmoles of SilencioM, dissolved in artificial CSF (aCSF), a buffer that mimics the electrolytes, osmolarity, and pH of native CSF (n=14). As controls, we injected mice with vehicle (aCSF) only (n= 6). We sacrificed all mice eight weeks after injection, intracardially perfused them with PBS and fixed one hemisphere for histology while the second was further dissected and flash frozen for biochemical studies (Figure 2b). To examine the cellular distribution of SilencioM, we employed *in situ* hybridization (ISH) using a Locked Nucleic Acid (LNA) modified probe that we designed against M2024, the guide strand. We found widespread cellular distribution of M2024 across the entire PS19 mouse brain (Figure 2c-e), confirming the uptake and retention of SilencioM for at least 2 months post injection.

**Figure 2.**
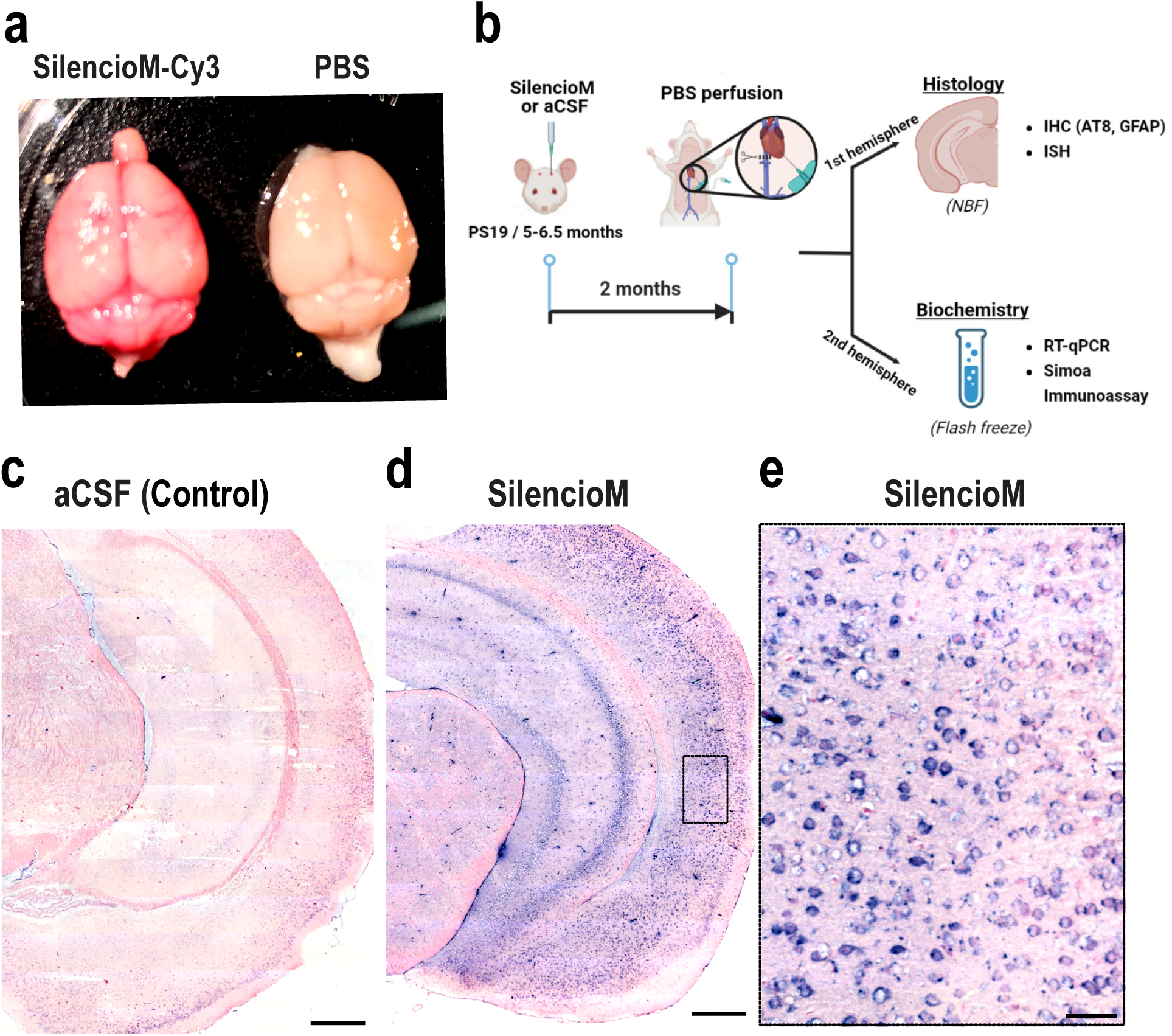
Widespread distribution and cell retention of SilencioM. **(a)** Gross brain images of B6 mice injected with SilencioM-Cy3 (red) or PBS, 2 days post injection. **(b)** Experimental workflow. **(c-e)** Representative images of LNA-ISH against M2024 in PS19 mouse brains. Widespread cellular uptake and distribution of M2024 (purple chromogen) is evident in SilencioM (**d, e**) but not in aCSF (**c**) injected PS19 mice; nuclear red = counterstain; scale bars: c, d = 500μm, e = 50μm. Panel b was created with BioRender.com

### SilencioM significantly reduces *MAPT* mRNA and tau in PS19 tauopathy mice

To assess functional target engagement by SilencioM, we developed an RT-qPCR assay for detection of human *MAPT* mRNA expressed from the transgene. We found significant reduction of human *MAPT* mRNA in all three brain areas of SilencioM injected mice compared to controls (posterior cortex 35.1%, p=0.007; hippocampus 37.2%, p=0.01; thalamus 20.4%, p=0.038) (Figure 3a; Supplementary Table 2).

**Figure 3.**
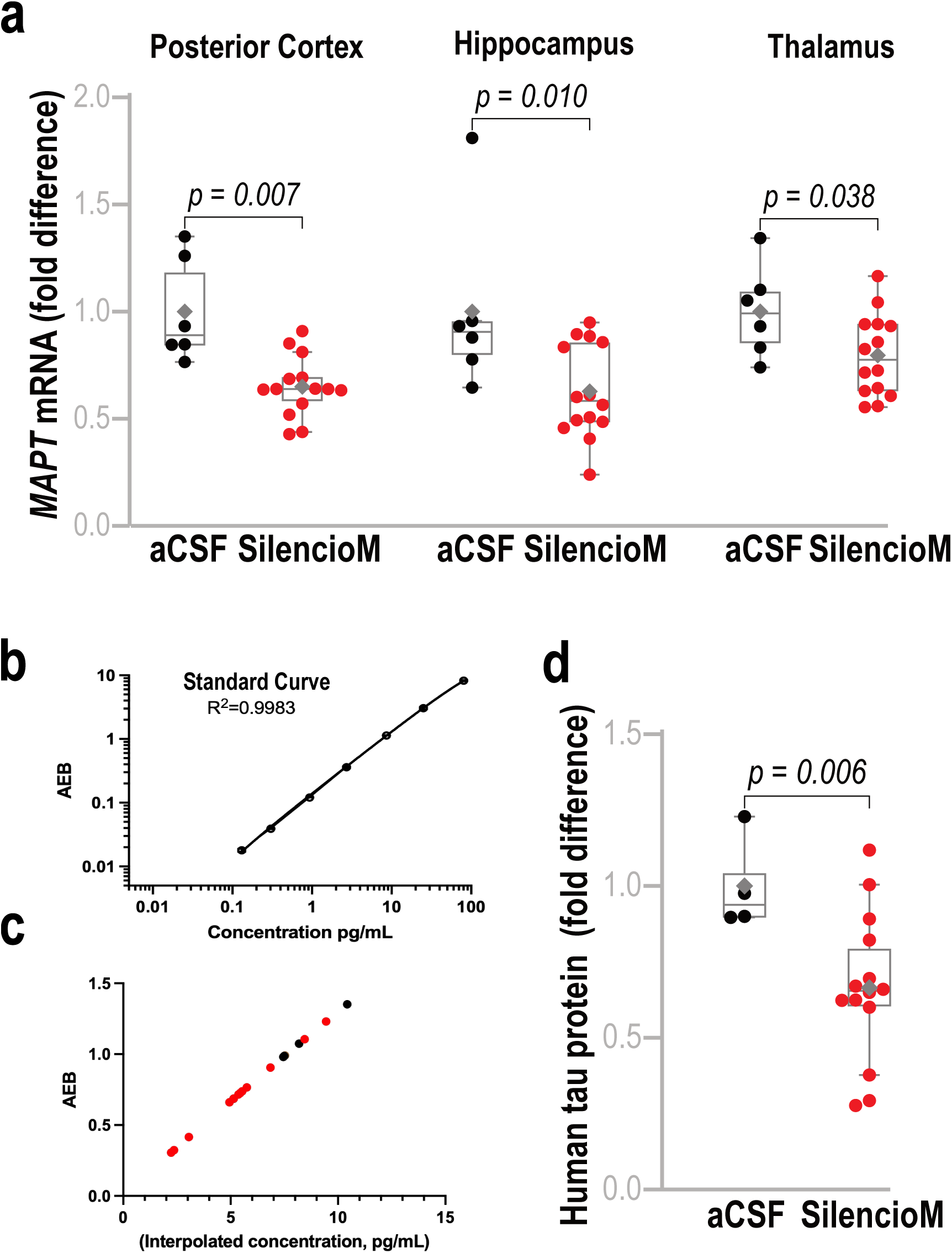
SilencioM significantly reduces *MAPT* mRNA and Tau protein in PS19 tauopathy mice. **(a)** Box plots of the fold difference of human *MAPT* mRNA in SilencioM (n=14) compared to aCSF (n=6) injected PS19 mice in indicated brain areas; each dot represents one mouse; median shown as horizontal grey line inside the box; mean is marked by a grey diamond. Posterior cortex and thalamus: unpaired, one-tailed, Welch’s t-test; hippocampus: unpaired, one-tailed, Mann-Whitney test. **(b-d)** Simoa immunoassay standardized for tissue lysates: (**b**) Standard curve generated from 8 samples containing known amounts of human tau. (**c**) Interpolated concentrations of human tau in the cortices of control (n=4, black dots) and SilencioM (n=14, red dots) injected PS19 mice. (**d**) Box plot of human tau concentration in cortices from aCSF and SilencioM injected PS19 mice; each dot represents a single mouse; unpaired, one-tailed, Welch’s t-test.

To examine the effect of SilencioM on human tau protein expressed from the transgene, we employed digital immunoassays based on the Single-Molecule Array (Simoa)-Quanterix platform. We established experimental parameters and dilutions to adjust Simoa for tissue samples, ran lysates from mouse posterior cortices of SilencioM and vehicle-only (aCSF) treated mice, and determined human tau concentration (Figure 3b, c). We found that SilencioM treatment caused a significant reduction (33,5%, p=0.006) of human tau (Figure 3d, Supplementary Table 3), indicating that SilencioM effectively targets *MAPT* for a durable reduction of tau.

### SilencioM effectively halts the development of tau pathology *in vivo*

PS19 mice show age-dependent aggregation of hyperphosphorylated tau (both NFTs and NTs), early synapse loss, astrogliosis and neuronal loss in advanced stages. To test the ability of SilencioM to affect the progression of tau neuropathology, we stained brain sections of treated and control PS19 mice (∼5.5 months of age at the time of injection) with AT8 antibody 2 months after injection. AT8 recognizes hyperphosphorylated tau (double epitope pSer202/pThr205) and is one of the most extensively characterized antibodies to detect NFTs, NTs and other forms of tau pathology [53] [35] [32]. We found that SilencioM treated mice showed substantial reduction in pathological tau throughout the entire brain compared to aCSF controls (81%, p=0.004) (Figure 4a-i; Supplementary Figure 2; Supplementary Table 4). This reduction was evident both in perikarya (NFTs) as well as in neuronal processes (NTs) (Figure 4e-h). To further examine the effect of SilencioM on astrogliosis, we stained astrocytes with glial fibrillary acidic protein (GFAP) and found significant reduction (26%, p=0.045), in astrogliosis upon treatment with SilencioM (Figure 5a-g, Supplementary Table 4). Taken together, our findings point to the efficacy of SilencioM in halting pathological tau development and, therefore, render it a promising candidate as a disease-modifying therapeutic targeting tauopathies including AD.

**Figure 4.**
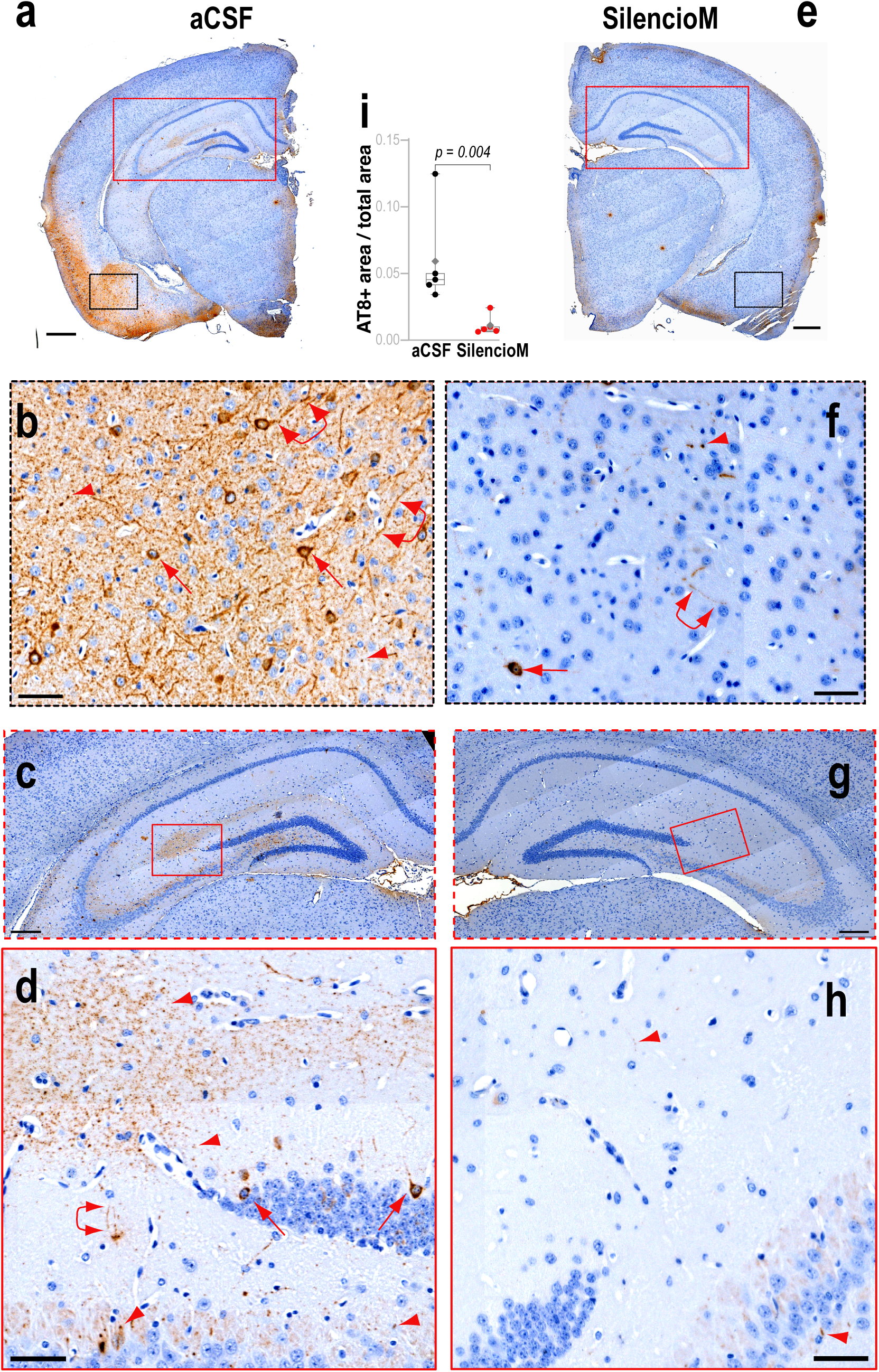
SilencioM ameliorates tau pathology *in vivo* Representative coronal hemispheric brain sections of PS19 mice injected with aCSF (**a-d**) or SilencioM (**e-h**), immunostained with AT8 (brown chromogen) and counterstained with hematoxylin (blue). Higher magnification images correspond to boxed areas from the hemisphere or image shown above. Numerous NFTs (arrows) and NTs (arrowheads: cross section; linked double arrowheads: longitudinal section) are evident in aCSF (**a-d**), while far fewer are seen in mice treated with SilencioM (**e-h**); scale bars: a, e= 500μm; b, f = 50μm; c, g = 200μm; d, h= 50μm. **(i)** AT8-positive area was quantified from matched coronal hemispheric sections of digitized slides from PS19 mice treated with SilencioM (n=5) or aCSF (n=5), normalized to total brain area per hemisphere, and plotted (each dot represents one mouse); one-tailed, unpaired, Mann-Whitney test.

**Figure 5.**
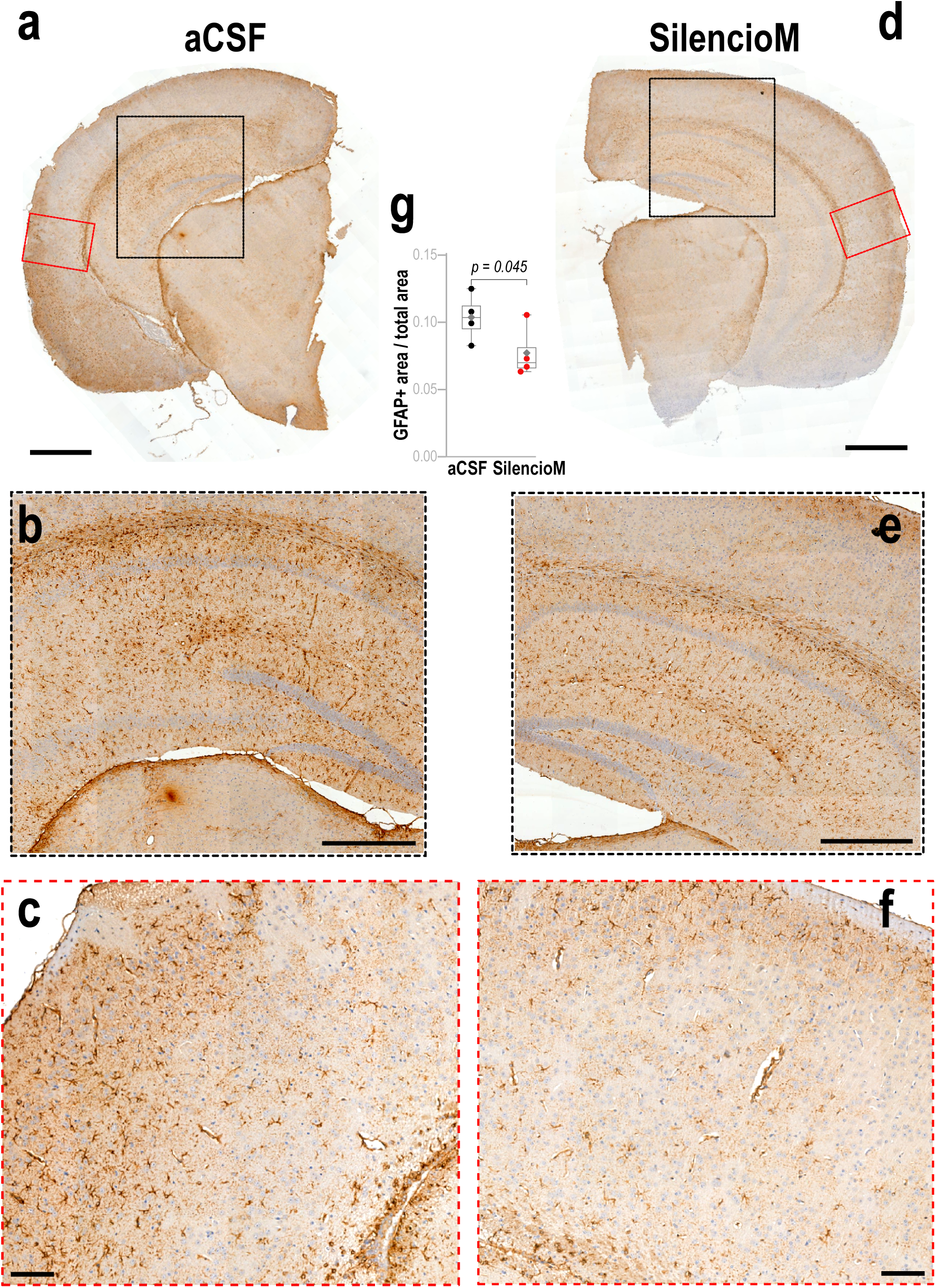
SilencioM ameliorates astrogliosis of tauopathy mice *in vivo* Representative coronal hemispheric brain sections of PS19 mice injected with aCSF (**a-c**) or SilencioM (**d-f**), immunostained for GFAP and counterstained with hematoxylin. Higher magnification images correspond to boxed areas from the hemisphere shown above. Decreased astrogliosis is evident in SilencioM (**d-f**) versus aCSF (**a-c**) treated PS19 mice; scale bars: a, d= 1000μm; b, e= 500μm: c, f= 100 μm. **(g)** GFAP-positive area was quantified from matched coronal hemispheric sections of digitized slides of PS19 mice treated with SilencioM (n=5) or aCSF (n=5), normalized to total brain area per hemisphere, and plotted (each dot represents one mouse); unpaired, one-tailed, Welch’s t-test.

## DISCUSSION

There is strong interest in developing therapeutics for tauopathies, given their prevalence and the lack of any treatments. Strategies that target tau are likely to have the greatest impact since tau is the proximal cause of disease pathogenesis. Here, we propose SilencioM as a new disease modifying therapeutic (DMT) for tauopathies, including AD. SilencioM targets a conserved region in human *MAPT* mRNA that is present in all isoforms and all major disease variants. As such, SilencioM should limit the production of all pathogenic tau protein isoforms (for example, 3Rtau and 4Rtau), conformers/fibrils and all oligomeric/soluble toxic species. We employed chemical modifications with known clinical safety data that enhance stability and reduce innate immune activation without abrogating Ago loading or silencing potency. SilencioM does not require encapsulation in LNPs or addition of lipophilic conjugates for delivery while being simple to manufacture and compatible with automated and large-scale synthesis under current Good Manufacturing Practices.

Reduction of tau protein levels in our experimental paradigm was accompanied by a significant decrease in tau pathology in PS19 mice two months after administration of the compound, which leaves open the possibility of SilencioM being active in suppressing pathological tau for even longer timeframes, thereby reducing the frequency of administration. Further studies are needed to optimize dose and administration frequency to achieve maximal pathology reduction with minimal side effects. Subsequent studies should also aim in exploring the therapeutic efficacy of SilencioM in older PS19 mice with established tau pathology. A limitation of our study, as with all preclinical studies, is the use of an animal model that cannot capture the full complexity of the human disease. Although PS19 mice are among the most widely used tauopathy models, they rely on overexpression of a human *MAPT* transgene with a pathogenic variant that causes early and severe tauopathy. As such, they develop rapid and severe pathology, unlike the slowly progressive nature of human tauopathies. Moreover, PS19 mice may not model the transmission and spreading of tau pathology. Future studies with wild-type mice seeded with patient-derived pathological tau are needed to examine the efficacy of SilencioM in targeting endogenous tau to halt the spread of pathology [42]. We anticipate that such studies, along with IND-enabling studies, can proceed smoothly since SilencioM was designed to target not only human *MAPT* but also primate, rat and mouse ortholog genes.

An advantage of siRNA-based designs is the ability to conjugate ligands to the passenger (sense) strand for targeted delivery, without disturbing the guide (antisense) strand, the active part of the molecule [52] [56] [26] [29]. Transferrin Receptor 1 (TfR1) binding and transcytosis is a particularly appealing strategy for shuttling therapeutics across the blood brain barrier after systemic delivery [43] [33] [34] [65] [71]. Many TfR1 ligands, including antibodies, nanobodies, peptides and aptamers are being developed, and the most mature designs have entered clinical trials [6] [7] [71] [58]. Conjugation of Tfr1 ligands in future studies will test SilencioM efficacy after subcutaneous injections and will expand the routes available for delivery. Importantly, the demonstration that the biological efficacy of the guide strand is retained when passenger strands are joined in a 5’ to 3’ manner, allows the development of multitargeting Silencios. In the accompanying manuscript we present SilencioMA, which simultaneously targets *MAPT* and *APP*, as a novel DMT specific for Alzheimer’s Disease.

## Conclusions

In this study we describe the design, development and testing of silencing connected interfering oligonucleotides against *MAPT* (SilencioM) as a potential novel DMT for tauopathies, including AD. SilencioM is effective in reducing the production of *MAPT* mRNA and tau protein in a mouse model of severe tauopathy after a single ICV injection. Importantly, SilencioM reduces tau pathology in the mouse brain for at least two months after administration. Because SilencioM is built on a modular scaffold, ligands that drive transcytosis across the blood-brain barrier can be attached, making systemic dosing possible. The straightforward production of SilencioM and its modular design make it a promising candidate for further studies.

## LIST OF ABBREVIATIONS

SilencioM: <u>Silen</u>cing <u>c</u>onnected interfering <u>o</u>ligonucleotides against *<u>M</u>APT*
MAPT: Microtubule Associated Protein Tau
siRNAs: Small interfering RNAs
AD: Alzheimer’s disease
IND: Investigational New Drug
mRNA: Messenger RNA
NFTs: Neurofibrillary tangles
NTs: Neuropil threads
VCP: Valosin-containing protein
PSP: Progressive Supranuclear Palsy
CBD: Corticobasal Degeneration
PiD: Pick’s disease
PART: Primary Age-related Tauopathy
APP: Amyloid Precursor Protein
ASOs: Antisense Oligonucleotides
CSF: Cerebrospinal fluid
ICV: Intracerebroventricular
LNPs: Lipid Nanoparticles
NIH: National Institutes of Health
NTC: Non-Targeting Control
Simoa: Single Molecule Array
ISH: In situ Hybridization
LNA: Locked Nucleic Acid
IHC: Immunohistochemistry
DIG: Digoxigenin
NBF: Neutral Buffered Formalin
GFAP: Glial fibrillary acidic protein
RT: Room temperature
BSA: Bovine Serum Albumin
UPLC: Ultra Performance Liquid Chromatography
MS: Mass Spectrometry
RLT: Recombinant human tau protein ladder
DMT: Disease Modifying Therapeutic
FDA: Food and Drug Administration

## ETHICS APPROVAL AND CONSENT TO PARTICIPATE

All procedures in this study were conducted in accordance with the National Institutes of Health (NIH) *Guide for the Care and Use of Laboratory Animals* and were reviewed and approved by the University of Pennsylvania Animal Care and Use Committee.

## CONSENT FOR PUBLICATION

Not applicable. This study does not contain any data from individual human subjects.

## AVAILABILITY OF DATA AND MATERIALS

All data generated or analyzed during this study are included in this published article (and its supplementary information files).

## AUTHORS’ CONTRIBUTIONS

Conceptualization: Z.M., Design and Methodology: Z.M., P. S., N.V. and H.X., Experimentation: P.S., N.V., K.X. and T.B., Data acquisition and curation: P.S. and N.V., Data analysis: P.S. and Z.M., Writing-original draft preparation: P.S., Writing-review and editing: P.S., Z.M., N.V., H.X., V.M-Y.L. and E.B.L., Project administration: P.S. and Z.M., Funding acquisition: Z.M. and E.B.L., Supervision: Z.M.

All authors read and approved the final manuscript.

## ACKNOWLEDGEMENTS

The authors would like to thank Dr. Kurt Brunden for many helpful discussions and comments on the manuscript, the Pathology Core Laboratory (RRID:SCR_028406) at the Children’s Hospital of Philadelphia Research Institute (Daniel Martinez) for providing brain sectioning services, and the Human Immunology Core (HIC, RRID: SCR_022380) at the Perelman School of Medicine at the University of Pennsylvania (Drs Yangzhu Du, Honghong Sun, and Eline Luning Prak) for assistance with Simoa assay.

---

**Supplementary Figure 1.**
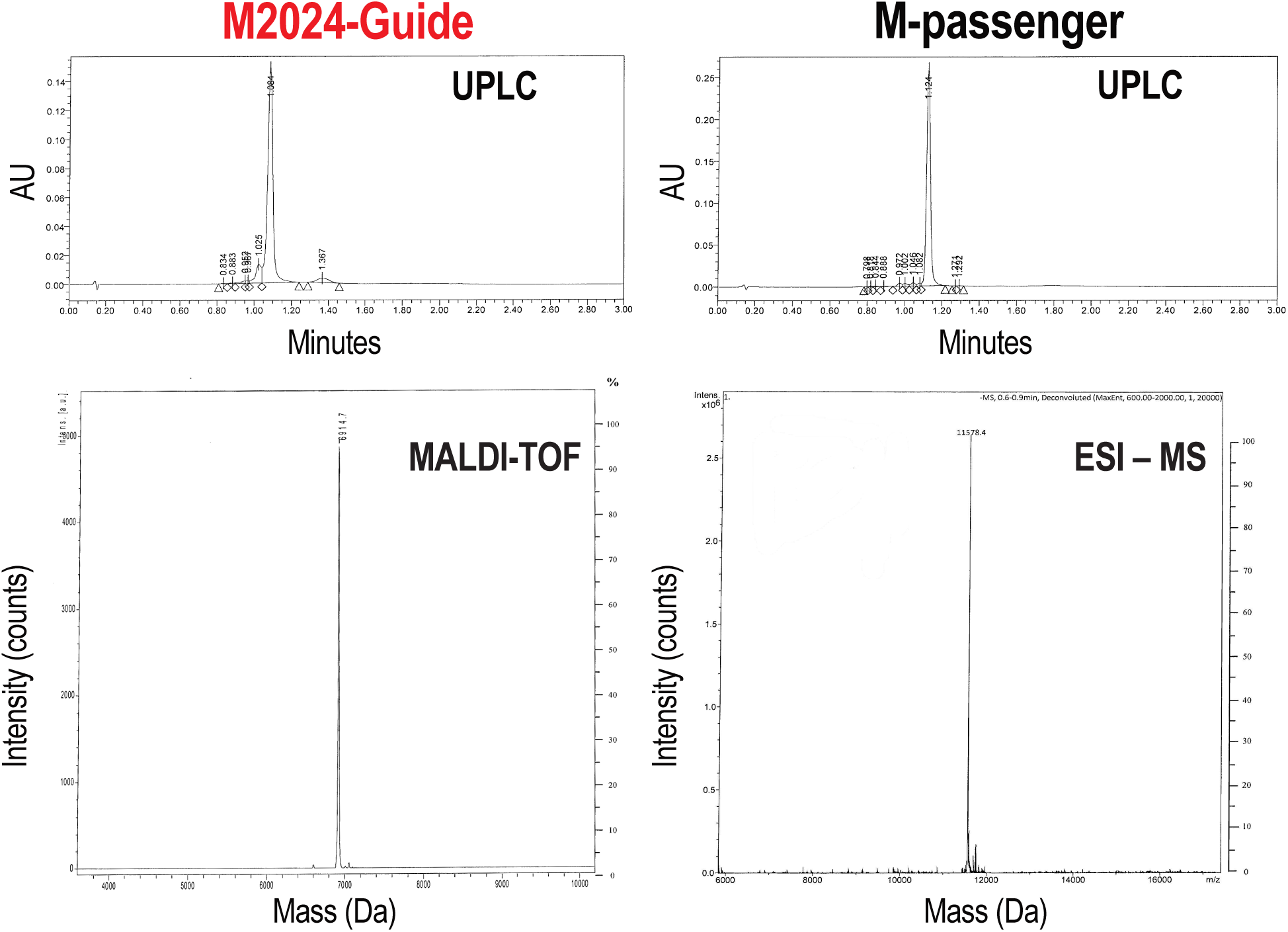
Properties of synthesized oligonucleotides. Ultra Performance Liquid Chromatography (UPLC) indicate the high purity of the oligonucleotides used to generate Silencios: M2024-Guide (85.9%), M-passenger (93.7%). Matrix Assisted Laser Desorption Ionization Time-Of-Flight (MALDI–TOF) or Electrospray Ionization Mass Spectrometry (ESI–MS) indicate that the synthesized oligonucleotides are of the expected masses: M2024-Guide (6912.7 Da theoretical, 6914.7 Da observed), M-passenger (11578.1 Da theoretical, 11578.4 Da observed).

**Supplementary Figure 2.**
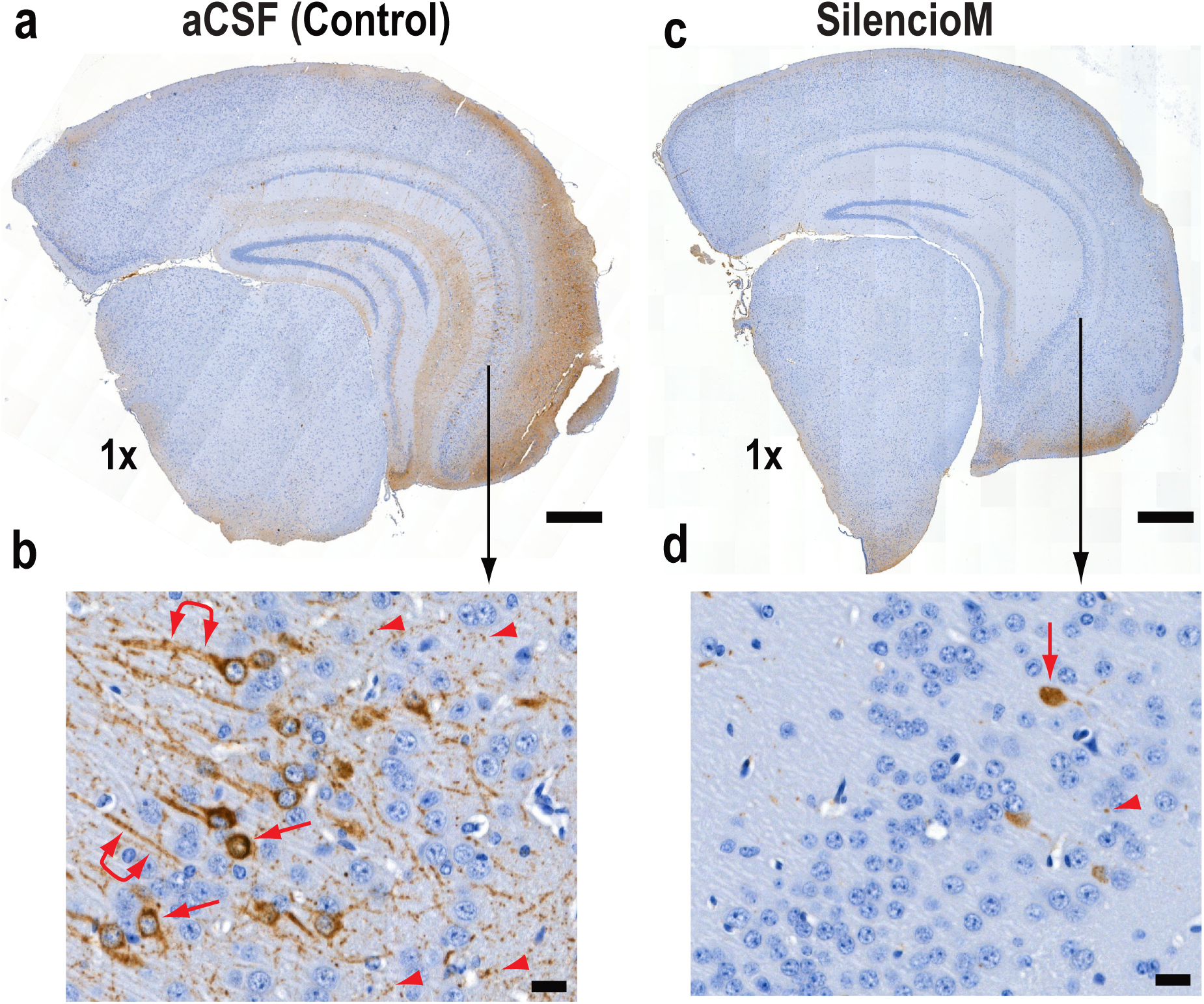
SilencioM ameliorates tau pathology *in vivo*. **(a-d)** Another example of AT8 staining in different PS19 mouse brains treated with aCSF (**a, b**) or SilencioM (**c, d**). Higher magnification images (**b, d**) correspond to areas, indicated by the black arrows, from the hemisphere shown above. Numerous NFTs (red arrows) and NTs (red arrowheads: cross section; linked red double arrowheads: longitudinal section) are evident in aCSF (**a, b**), while they are substantially reduced in mice treated with SilencioM (**c, d**); scale bar in a, c = 500 microns; b, d = 20 microns.

