## Supplementary Table 1 for "SilencioM: silencing connected interfering oligonucleotides against *MAPT* as a potential disease modifying therapeutic for tauopathies"

| Name | 5´-3´sequence | Target |
|---|---|---|
| MAPT-F | CTTAGCAACGTCCAGTCCAA | MAPT in HEK293 cells |
| MAPT-R | TCACCTTGCTCAGGTCAAC | |
| TgMAPT-F | GAAGATTGGGTCCCTGGAC | MAPT human transgene in PS19 |
| TgMAPT-R | GTCTGTCTTGGCTTTGGCG | |
| HsGAPDH-F | GAGTCCACTGGCGTCTTCAC | GAPDH in HEK 293 cells |
| HsGAPDH-R | GAGGCATTGCTGATGATCTTGAG | |
| MmGAPDH-F | CATCACTGCCACCCAGAAGACTG | GAPDH in PS19 mice |
| MmGAPDH-R | ATGCCAGTGAGCTTCCCGTTCAG | |
