## Supplementary Table 2 for "SilencioM: silencing connected interfering oligonucleotides against *MAPT* as a potential disease modifying therapeutic for tauopathies"

| Sample Name | Group | Target Name | Posterior cortex Ct mean | Hippocampus Ct mean | Thalamus Ct mean |
|---|---|---|---|---|---|
| 1 | Control (aCSF) | GAPDH | 17.81 | 16.91 | 17.48 |
| 1 | Control (aCSF) | MAPT | 18.62 | 17.91 | 19.80 |
| 2 | Control (aCSF) | GAPDH | 17.45 | 18.97 | 17.43 |
| 2 | Control (aCSF) | MAPT | 18.39 | 19.93 | 19.60 |
| 3 | Control (aCSF) | GAPDH | 17.61 | 21.93 | 17.39 |
| 3 | Control (aCSF) | MAPT | 18.55 | 21.97 | 19.89 |
| 4 | Control (aCSF) | GAPDH | 17.57 | 18.59 | 17.51 |
| 4 | Control (aCSF) | MAPT | 17.85 | 19.85 | 19.15 |
| 5 | Treated (SilencioM) | GAPDH | 17.40 | 18.18 | 17.50 |
| 5 | Treated (SilencioM) | MAPT | 18.41 | 19.30 | 20.41 |
| 6 | Treated (SilencioM) | GAPDH | 17.47 | 17.70 | 17.60 |
| 6 | Treated (SilencioM) | MAPT | 19.40 | 18.77 | 20.30 |
| 7 | Treated (SilencioM) | GAPDH | 17.54 | 17.22 | 17.54 |
| 7 | Treated (SilencioM) | MAPT | 18.89 | 18.94 | 20.27 |
| 8 | Treated (SilencioM) | GAPDH | 17.29 | 17.42 | 17.77 |
| 8 | Treated (SilencioM) | MAPT | 18.65 | 19.30 | 20.29 |
| 9 | Treated (SilencioM) | GAPDH | 17.61 | 17.19 | 17.41 |
| 9 | Treated (SilencioM) | MAPT | 18.45 | 18.80 | 19.57 |
| 10 | Treated (SilencioM) | GAPDH | 17.71 | 17.51 | 17.80 |
| 10 | Treated (SilencioM) | MAPT | 19.61 | 19.44 | 20.70 |
| 11 | Treated (SilencioM) | GAPDH | 17.62 | 17.39 | 17.59 |
| 11 | Treated (SilencioM) | MAPT | 18.98 | 19.02 | 20.37 |
| 12 | Treated (SilencioM) | GAPDH | 17.27 | 19.83 | 17.78 |
| 12 | Treated (SilencioM) | MAPT | 18.63 | 20.99 | 19.78 |
| 13 | Treated (SilencioM) | GAPDH | 16.56 | 25.35 | 17.66 |
| 13 | Treated (SilencioM) | MAPT | 17.79 | 27.55 | 19.81 |
| 14 | Treated (SilencioM) | GAPDH | 17.02 | 24.77 | 17.21 |
| 14 | Treated (SilencioM) | MAPT | 17.96 | 26.80 | 19.05 |
| 15 | Treated (SilencioM) | GAPDH | 16.72 | 19.10 | 17.73 |
| 15 | Treated (SilencioM) | MAPT | 18.37 | 22.06 | 20.27 |
| 16 | Treated (SilencioM) | GAPDH | 16.84 | 18.66 | 17.92 |
| 16 | Treated (SilencioM) | MAPT | 18.19 | 19.71 | 20.26 |
| 17 | Treated (SilencioM) | GAPDH | 16.90 | 19.19 | 17.43 |
| 17 | Treated (SilencioM) | MAPT | 18.15 | 20.17 | 19.58 |
| 18 | Treated (SilencioM) | GAPDH | 16.83 | 18.82 | 17.46 |
| 18 | Treated (SilencioM) | MAPT | 18.34 | 20.73 | 19.74 |
| 19 | Control (aCSF) | GAPDH | 16.70 | 18.72 | 17.32 |
| 19 | Control (aCSF) | MAPT | 17.79 | 20.24 | 19.24 |
| 20 | Control (aCSF) | GAPDH | 18.45 | 18.98 | 17.72 |
| 20 | Control (aCSF) | MAPT | 18.83 | 20.06 | 19.71 |
