## Supplementary Table 3 for "SilencioM: silencing connected interfering oligonucleotides against *MAPT* as a potential disease modifying therapeutic for tauopathies"

| Sample ID | Plex | Mean AEB | SD Conc. | CV Conc. | Mean Conc. | Treatment |
|---|---|---|---|---|---|---|
| PC1 | Tau | 0.034 | 0.154 | 14% | 1.074 | aCSF |
| PC3 | Tau | 0.032 | 0.099 | 10% | 0.991 | aCSF |
| PC4 | Tau | 0.042 | 0.026 | 2% | 1.352 | aCSF |
| PC5 | Tau | 0.035 | 0.039 | 4% | 1.106 | SilencioM |
| PC6 | Tau | 0.025 | 0.094 | 13% | 0.715 | SilencioM |
| PC7 | Tau | 0.024 | 0.015 | 2% | 0.686 | SilencioM |
| PC8 | Tau | 0.026 | 0.098 | 13% | 0.765 | SilencioM |
| PC9 | Tau | 0.024 | 0.013 | 2% | 0.687 | SilencioM |
| PC10 | Tau | 0.014 | 0.011 | 3% | 0.323 | SilencioM |
| PC11 | Tau | 0.030 | 0.064 | 7% | 0.905 | SilencioM |
| PC12 | Tau | 0.025 | 0.020 | 3% | 0.726 | SilencioM |
| PC13 | Tau | 0.039 | 0.029 | 2% | 1.231 | SilencioM |
| PC14 | Tau | 0.032 | 0.007 | 1% | 0.981 | SilencioM |
| PC15 | Tau | 0.017 | 0.006 | 2% | 0.415 | SilencioM |
| PC16 | Tau | 0.023 | 0.051 | 8% | 0.661 | SilencioM |
| PC17 | Tau | 0.025 | 0.049 | 7% | 0.738 | SilencioM |
| PC18 | Tau | 0.014 | 0.048 | 16% | 0.305 | SilencioM |
| PC19 | Tau | 0.032 | 0.061 | 6% | 0.987 | aCSF |
