## Supplementary Table 4 for "SilencioM: silencing connected interfering oligonucleotides against *MAPT* as a potential disease modifying therapeutic for tauopathies"

| Total area (µm^2) | AT8: Positive area (µm^2) | Treatment |
|---|---|---|
| 44260861.7 | 1513604.4 | aCSF |
| 48970267.2 | 2223695.2 | aCSF |
| 29519247.6 | 3683236 | aCSF |
| 29682908.7 | 1232146.75 | aCSF |
| 42061148.2 | 2102208.5 | aCSF |
| 33609826.6 | 212203.4 | SilencioM |
| 31077306.1 | 215588.1 | SilencioM |
| 31941338.1 | 318289.3 | SilencioM |
| 34270962.1 | 280179.2188 | SilencioM |
| 32948863.5 | 799767.4375 | SilencioM |

| Total area (µm^2) | GFAP: Positive area (µn | Treatment |
|---|---|---|
| 31523325.3 | 2604215.2 | aCSF |
| 41786931.6 | 5221026 | aCSF |
| 22472332.5 | 2230395.8 | aCSF |
| 27347335.2 | 2950596 | aCSF |
| 35326852.5 | 2365468 | SilencioM |
| 29990075.4 | 3162671.5 | SilencioM |
| 26146065.1 | 1655872.875 | SilencioM |
| 31055204.9 | 2266659.8 | SilencioM |
